# Biophysical characterization of novel biomarkers and bioaffinity reagents (NanoMIPs): on the road for a low-cost diagnostic test for intestinal schistosomiasis

**DOI:** 10.64898/2026.07.28.741194

**Authors:** Flavio Isopo, Andrei Nino Stephen, Giovanna Boumis, Alessandra Giorgi, Ivano Eberini, Subrayal M. Reddy, Adriana Erica Miele

**Affiliations:** Université Lyon 1, CNRS, UMR 5280 – Institut des Sciences Analytiques, 5 Rue de la Doua, 69100 Villeurbanne, France; Centre for Smart Materials, University of Lancashire, Preston, PR1 2HE, UK; Dept. Biochemical Sciences, Sapienza University of Rome, P.le Aldo Moro 5, 00185 Rome, Italy; Dipartimento di Scienze Farmacologiche e Biomolecolari “Rodolfo Paoletti”, Università degli Studi di Milano, via Giuseppe Balzaretti 9, 20133 Milano, Italy

**Keywords:** SmVAL, Molecularly Imprinted Polymers, SAXS, Molecular Modeling, biolayer interferometry BLI, Electrochemical Impedance Spectroscopy, Neglected Tropical Diseases, Biomarker, Diagnostic

## Abstract

*Schistosoma mansoni* is a vector-borne intestinal parasite, endemic in ∼70 tropical countries. Most of the World Health Organization interventions are based on mass drug administration (MDA) and sanitation. Since the parasite does not induce permanent immunity, reinfection rate is high, inducing scheduling of several MDA campaigns. To follow up after treatment and avoid blind MDA, providing a new rapid and cheap diagnostic test is needed, since the most used method is fifty years-old, lacks sensitivity and efficiency. Venom Allergen-Like proteins (SmVALs) have been previously identified as secreted/excreted proteins and potential biomarkers. Here we present the biophysical characterization of SmVAL11 and SmVAL13, their recognition by both specific antibodies and molecularly imprinted polymers (nanoMIPs), an easy-to-standardize alternative. The specific antibodies showed no cross-reactivity, while initial characterization of the nanoMIPs indicates a promising level of selectivity, establishing them as a reliable, cost-effective alternative. Therefore, our results are promising for the future development of a new cheap diagnostic test of intestinal schistosomiasis.

## Introduction

Schistosomiasis is a vector-borne parasitic disease, caused by trematodes from the *Schistosoma* genus, with a highly debilitating chronic onset and a low mortality rate [McManus et al., 2018]. At present it is endemic in about 70 countries, but due to global warming it is spreading over the tropics, reaching, for example, the mediterranean area [Berry et al., 2016] [Salas-Coronas et al., 2021]. The disease does not induce permanent humoral immunity; hence the reinfection rate is high, particularly in endemic countries. The World Health Organization (WHO) has launched several campaigns of Mass Drug Administration (MDA), with a focus on school-age children and adults [King et al., 2019], with the aim of reducing the burden and eliminate most neglected tropical diseases (NTD) by 2030, as per the RoadMap 2030 [WHO, 2017].

The gold-standard drug for MDA is Praziquantel, a multi-target molecule [Thomas et al., 2018; Angelucci et al., 2007] to which resistance has not yet arisen, although some strains with reduced sensitivity have been reported [Doenhoff et al., 2002; Sisay et al., 2025]. Diagnostic-wise, WHO guidelines recommend the Kato-Katz test, which is based on the microscopic examination of stools and urine after microfiltration and concentration, to find eggs and therefore active infection [WHO, 2019]. Even though this test is cost-effective and does not need particular infrastructures, it lacks sensitivity and cannot be considered as an early diagnostic procedure, making it less effective in decreasing the burden of the parasite. In a few countries this test is complemented with a handful of more sensitive tests based on the detection of genetic material or of circulating sugar antigens [Silva-Moraes et al., 2019; Casacuberta-Partal et al., 2019]. Although more sensitive, the latter’s half-life is longer than the infection itself, making follow-up diagnosis unreliable. The sensitivity of the PCR based method is higher, but its cost is still not affordable in many endemic countries, and it needs both trained healthcare workers and expensive equipment, consumables and reagents, the latter requiring cold storage. Thus, to avoid blind MDA, to prevent a rise in resistance and to advance in the field of diagnosis, a rapid and cheap diagnostic test needs to be produced and delivered.

In the recent past, Chalmers and Farias [Farias et al., 2019; Chalmers et al., 2008] discovered a life-cycle related expression pattern of a high variable family of proteins in *S. mansoni*, the Venom Allergen-Like (SmVAL) proteins. SmVALs are a family of at least 29 isoforms [Farias et al., 2019] and are part of the superfamily of cysteine-rich secretory proteins, antigen 5 and pathogenesis-related 1 (CAP/PR-1) proteins, which is subdivided in the family of cysteine-rich secretory proteins (CRISP) and in the family of the Sperm-Coating Protein/ Tpx/ Antigen5/ Pathogenesis-related-1/ Sc7 (SCP/TAPS). This latter structural domain is found in many species, including humans and plants, over and above trematodes. The functions of the SCP/TAPS domain ranges from immunologic regulation to crosstalk and development, though their precise role in schistosomes is currently unknown [Cantacessi et al., 2012].

Despite SmVALs belonging to the SCP/TAPS family, they display low sequence identity with the other family members (19% on average). However, the identity is high (70-89%) within the 29 SmVALs [Chalmers et al., 2008]. Sequence-wise SmVALs are subdivided into two subgroups: Group 1 is composed by SmVAL 1-5, 7-10, 12, 14, 15, 18-29 while group 2 includes SmVAL 6, 11, 13, 16 and 17. Group 1 harbors a signal peptide and contains 6 conserved Cys residues; whereas group 2 has not a signal peptide and contains only 4 Cys. [Chalmers et al., 2008]. To the best of our knowledge, the structure of only one isoform, SmVAL4, has been solved by X-ray crystallography (PDB: 4P27, [Kelleher et al., 2014]), leaving the need to solve the structure of the other isoforms and understand their function and life-cycle regulation.

Given the different SmVALs isoforms, it is crucial to select the most suitable candidate for a diagnostic device development. For this purpose, we have chosen SmVAL13 and SmVAL11, two isoforms that exhibit high expression levels at distinct stages of the parasite’s life cycle: early (juvenile and female worms) for smVAL13 and late (eggs, adult male and female worms) for SmVAL11 [Chalmers et al., 2008].

While biological antibodies have been developed against both isoforms, their high production costs limit large-scale deployment for protein detection, thereby constraining efforts to track and treat *S. mansoni*, which is responsible for their expression. This challenge is further exacerbated by the *S. mansoni* prevalence in resource-limited regions, including sub-Saharan Africa, parts of South America, the Caribbean, and the Middle East [Olveda, 2013], where reliable refrigeration infrastructure and cold-chain logistics, which are essential for antibody storage and use are often lacking [Talbot, de Koning-Ward, & Layton, 2025]. Consequently, antibody-based diagnostics can be unreliable or impractical in these settings, emphasizing the need for more robust alternatives. Molecularly imprinted polymers (MIPs) are synthetic, polymeric bio-affinity reagents that offer a promising solution. As chemically produced materials, MIPs exhibit high stability across a wide range of conditions, including at and above room temperature [Basak, Venkatram, & Singhal, 2022], eliminating the need for cold-chain storage. They have been extensively investigated for a variety of targets, including small molecules [Farooq et al., 2018], peptides [Herrera León et al., 2023; Hoshino et al., 2008], proteins [Reddy et al., 2024; Sullivan, Clay, et al., 2021], and viruses [El Sharif et al., 2022]. MIPs can be synthesized rapidly (typically within 24 hours) in a one-pot process using low-cost, rationally selected functional monomers [Sullivan et al., 2019] that self-assemble around a target template molecule. Following cross-linked polymerization, removal of the template yields specific binding sites (imprints), capable of high-affinity and selective rebinding of the target [Stephen & Reddy, 2025]. Recent advances in bottom-up synthesis approaches have enabled the development of nanoscale MIPs (nanoMIPs), which demonstrate superior binding affinity and selectivity for protein targets, in some cases surpassing that of animal-derived antibodies [Smolinska-Kempisty et al., 2016; Stephen, Boldrin, & Reddy, 2026], positioning them as highly attractive, animal-free alternatives for immune-diagnostic applications [Kaur et al., 2024; Stephen & Reddy, 2025]. Altogether these characteristics make nanoMIPs an ideal low-cost alternative that can be rapidly manufactured and deployed in resource-constrained settings.

In this work we present the biophysical characterization of the recombinant forms of SmVAL11 and SmVAL13 together with the analysis of the complexes with their cognate polyclonal antibodies (pAb), and nanoMIPs with the aim to use them as biomarkers for early schistosomiasis detection and treatment follow-up.

## Results

### Cloning, expression and production of recombinant SmVAL13 and SmVAL11

SmVAL13 and SmVAL11 were cloned into the pTXB1 vector, containing at the C-Terminus an autocleavable Intein Tag and a Chitin Binding Domain (CBD). This tag avoids using expensive proteases and does not leave extra amino acids after the cleavage. Affinity purification was carried out on a Chitin Resin and elution, concomitant with tag elimination, was done with high salt buffer containing ditiothreitol (DTT). Final yields that were obtained were at least 10 mg/L of culture.

### SmVAL13 biophysical characterization

SmVAL13 is a 27.2 kDa and basic protein (Table 1) that contains one single SCP/TAPS domain, followed by a C-terminal long and flexible tail. After purification, SDS-PAGE at 15% was done to assess the purity of the sample in denaturing conditions, the result (Fig. 1a) revealed a prominent band at around 30 kDa, a faint one at 60 kDa and no apparent degradation. Western Blot was performed to rule out the presence of contaminants. As shown in Fig. 1a, pAb anti-SmVAL13 recognizes both the band at 30 kDa and at 60 kDa, suggesting a possible dimerization through the Cys residues.

**Fig. 1.**
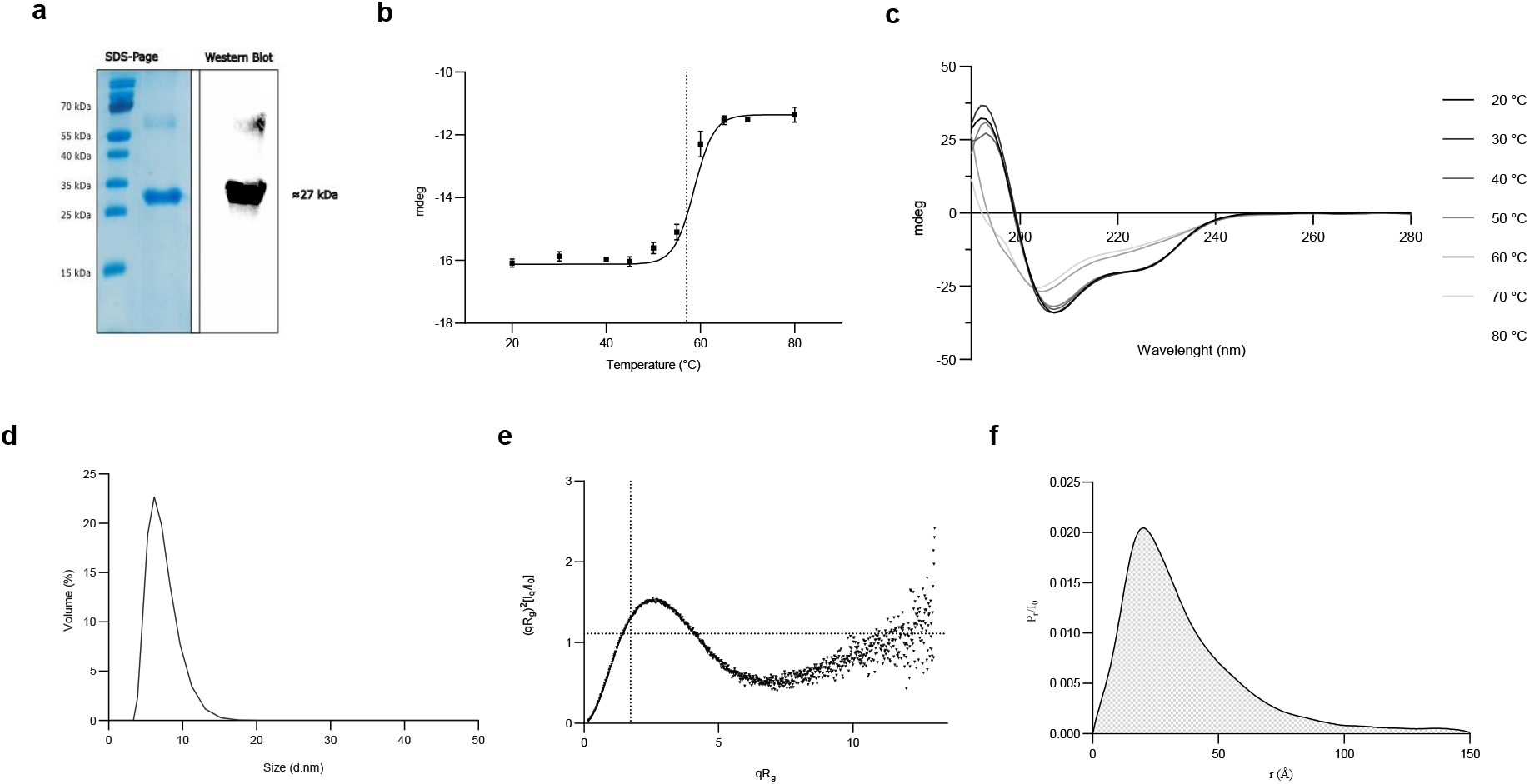
Summary of SmVAL13 data analysis. (a) SDS-PAGE at 15% and Western Blot; (b) Circular Dichroism thermal ramping spectra (from 20 to 80 °C) of SmVAL13 (10 μM) in 2.5 mM MES pH 6.8, 12.5 mM NaCl, 0.1 mM EDTA, each CD spectrum is the average of 5 spectra after subtraction of the averaged blank (buffer only) and smoothed with Savitzky-Golay over 3 points; (c) ellipticity (mdeg) at 222 nm plotted against temperature ramping, Boltzmann fit curve in black, experimental points with standard errors (N=3) in black squares; (d) extrapolation of Rh from Dynamic Light Scattering data, of SmVAL13 (50 μM) in 25 mM MES pH 6.8, 125 mM NaCl, 1 mM EDTA; (e) Normalized Kratky plot of SAXS data (extrapolated to zero concentration); (f) Pr function of data in (e).

**Table 1.** Summary of some characteristics of SmVALs characterized in this paper.

| Isoform | MW (from gene) | Theoretical pI [Gasteiger et al., 2005] | Parasitic stage of maximal mRNA expression (qRT-PCR) [Chalmers et al., 2008] |
| --- | --- | --- | --- |
| SmVAL11 | 45.0 kDa | 8.3 | Eggs, adult male and female worms |
| SmVAL13 | 27.2 kDa | 9.5 | Juvenile and adult female worms |

Further quality control check was performed with dynamic light scattering (DLS) and circular dichroism (CD). In the first case one single peak centered at a hydrodynamic radius (R_h_) of 6.1 nm is found, compatible with a mono-dispersed, yet elongated monomeric protein (Fig. 1d). CD spectra indicate a well folded protein, with a mixed secondary structure of α-helices and β-strands, compatible with the SCP/TAPS fold (Fig. 1b). Thermal denaturation of purified sample indicates substantial stability, as the denaturation remains incomplete up to 80 °C, however the molten-globule-like transition appears to be irreversible (data not shown). Overall, the calculated apparent Tm is 57 ± 2.5 °C (Fig. 1c), assuming a 2-states transition.

In order to gain insight into the tertiary structure, we performed small angle scattering (SAXS) with synchrotron light, in a dilution series starting at 7 mg/ml. The normalized Kratky plot (Fig.1e) is indicative of a globular protein with an elongated part, in fact SmVAL13 deviates from the bell-shaped profile, suggesting the presence of flexible parts, most probably the C-terminal extension, which is predicted to be low complexity/intrinsically disordered. This behavior is also evident from the P(r) pair-distribution function (Fig.1f), which shows a non-symmetrical peak ending at D_max_ = 15.1 nm. The ratio D_max_/R_g_ = 5.8 deviates from a perfect spherical particle. Moreover, the extracted molecular weight of 21.8 kDa (Bayesian inference) is into agreement with the theoretical mass of SmVAL13 (27.2 kDa) and further indicates the monomeric assembly of the protein in solution.

### SmVAL11 biophysical characterization

SmVAL11 is a 45 kDa basic protein (Table 1), which contains two SCP/TAPS domains in tandem linked by a flexible linker, possibly originating from a gene duplication event [Chalmers et al., 2008]. As shown in figure 2a, SDS-PAGE revealed multiple bands, one at 110 kDa, one at 48 kDa and three others below 20 kDa. MALDI Mass Spectrometry analysis was performed after trypsin digestion of the protein extracted from the bands (Fig. 2b, species A, B, C and D), which highlighted the cleavage of SmVAL11. Despite this apparent dis-homogeneity, DLS (Fig.2c) identified only one population, with a Rh of ∼5.3 nm, suggesting a possible interaction in solution between the different SmVAL11 populations present in the SDS-PAGE.

**Fig. 2.**
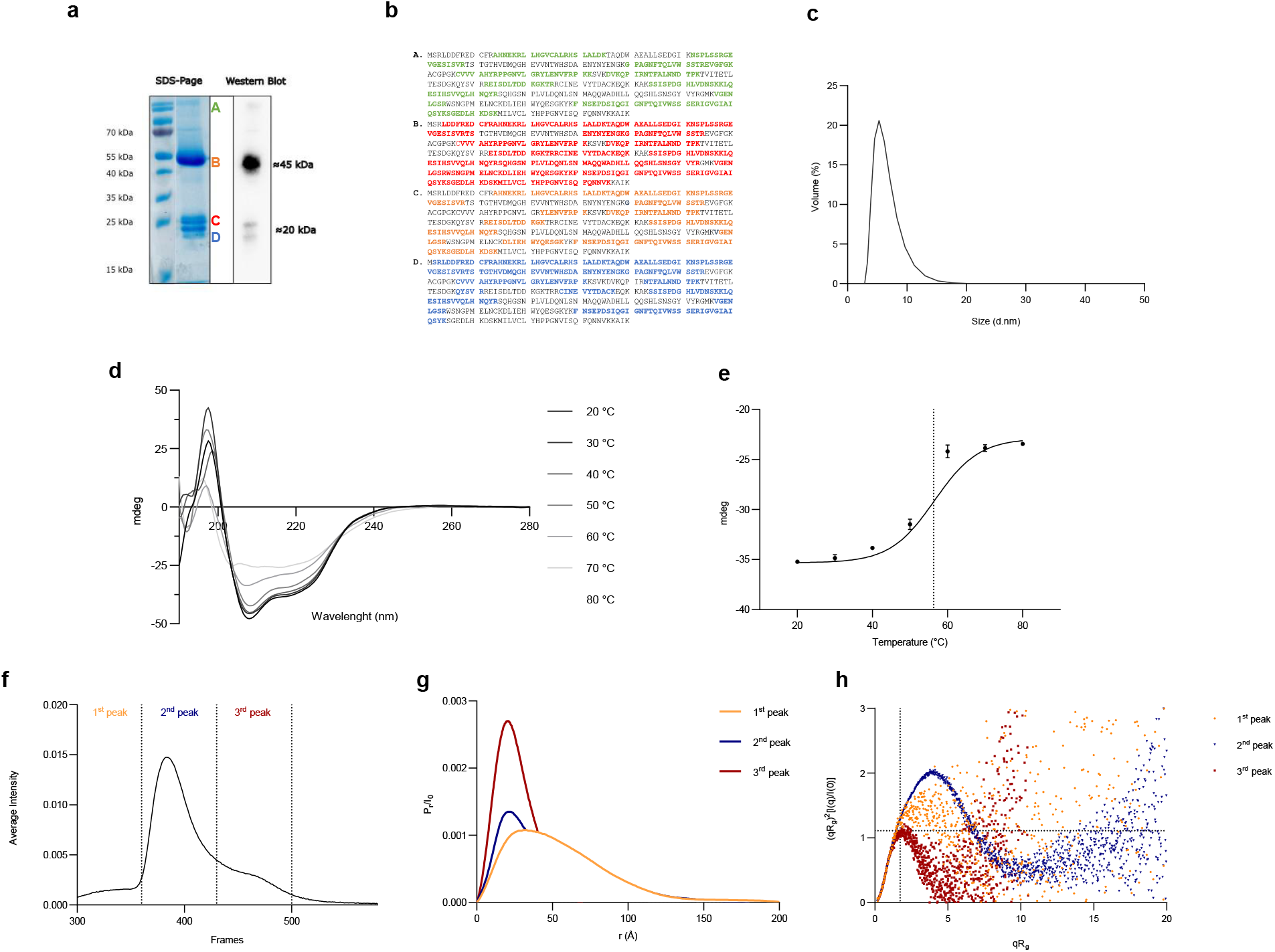
Summary of SmVAL11 data analysis. (a) SDS-PAGE at 15% and Western Blot; (b) MALDI analysis of different populations of SmVAL11 after trypsin digestion; (c) extrapolation of Rh from Dynamic Light Scattering data of SmVAL11 (50 μM) in 25 mM MES pH 6.8, 125 mM NaCl, 1 mM EDTA; (d) Circular Dichroism thermal ramping spectra (from 20 to 80 °C) of SmVAL11 (10 μM) in 2.5 mM MES pH 6.8, 12.5 mM NaCl, 0.1 mM EDTA; each CD spectrum is the average of 5 spectra after subtraction of the averaged blank (buffer only) and smoothed with Savitzky-Golay over 3 points; (e) ellipticity (mdeg) at 222 nm plotted against temperature ramping, Boltzmann fit curve in black, experimental points with standard errors (N=3) in red; (f) SEC-SAXS chromatogram of SmVAL11; (g) superposed Pr function of the three eluted peaks after deconvolution; (h) Normalized Kratky plot of the three eluted peaks, after deconvolution with EFA.

Similarly to SmVAL13, SmVAL11 CD spectra at 20 °C (Fig. 2d) are characteristic of a well folded protein, with a mix of α-helices and β-strands, again consistent with the SCP/TAPS domain. Thermal denaturation behavior was also very similar between the two proteins (Fig. 2e) with an apparent T_m_ of 54 ± 1 °C, and an incomplete though irreversible unfolding (data not shown).

SAXS measurements coupled to Size Exclusion Chromatography (SEC-SAXS) were then performed to separate and analyze the different populations. As shown in Fig. 2f, the integrated intensity chromatogram gave a partial separation of three populations identified as three partially overlapping peaks. Through Evolving Factor Analysis (EFA, Hopkins JB. 2024) the peaks were deconvoluted and analyzed separately. The first peak represents a minor population of large particles (Fig. 2g) possibly composed by several SmVAL11 (Table 2). The second peak presents the features of a flexible multi-domain particle (Fig. 2h) with R_g_ = 3.87 nm and D_max_ = 18 nm, calculated from the P_(r)_ function. The calculated molecular weight is 40.2 kDa (Table 2), in agreement with the theoretical MW of SmVAL11 (45 kDa). The third peak is a compact globular species of about 18 kDa, compatible with the size of one auto-cleaved SCP/TAPS domains.

**Table 2.** Summary of SEC-SAXS data analysis after deconvolution with EFA for SmVAL11. R_g_ and D_max_ are calculated with GNOM and MW through Bayesian inference.

| SmVAL11 | $R_g$ | $D_{max}$ | MW |
| --- | --- | --- | --- |
| 1<sup>st</sup> Peak | $4.21 \pm 0.05$ nm | 20.4 nm | 74.3 kDa |
| 2<sup>nd</sup> Peak | $3.87 \pm 0.01$ nm | 18 nm | 40.2 kDa |
| 3<sup>rd</sup> Peak | $1.90 \pm 0.01$ nm | 7 nm | 18.1 kDa |

### SmVAL11 and SmVAL13 Antigenicity and homology structures

Having established that both proteins are stable and well-folded, we assessed their suitability as isoform-specific serological targets. Linear B-cell epitopes were predicted with BepiPred [Jespersen et al., 2017] and mapped onto the secondary-structure and into disorder profiles obtained from PSIPRED and DISOPRED [Jones, 1999; Jones & Cozzetto, 2015], allowing antigenicity to be correlated with local structure (Fig. 3a, 3b). For SmVAL13, the highest epitope scores cluster from residue 160 onward, overlapping with the disordered C-terminal tail (Fig. 3a); for SmVAL11, the dominant antigenic signal localizes to the inter-domain linker (Fig. 3b). In both proteins the strongest epitopes map to coil and flexible regions, the segments most accessible to antibody binding. Since sequences of these regions are divergent between the two isoforms, antibodies raised against each protein are predicted not to cross-react. To rationalize this specificity at the structural level, and to exclude cross-reactivity with the host proteome, we searched for homologues by BLAST against UniProtKB [UniProt Consortium, 2025]. No human protein showed detectable similarity to SmVAL11, and only four human CAP-family members shared ≥60% similarity with SmVAL13: GLIPR1L1 (Q6UWM5, sperm-associated), GLIPR1 (P48060, glioma-related), GLIPR2/GAPR-1 (Q9H4G4, brain-associated) and PI16 (Q6UXB8, tumor-associated). All four are either non-circulating or confined to brain, sperm or tumor tissue and are absent or minimally expressed in blood under physiological conditions, making cross-recognition in blood negligible. We then built homology models of SmVAL13 and of the two SmVAL11 domains (SmVAL11-N and SmVAL11-C) independently (Fig. 3c) and superposed all three onto their closest templates (4AIW and 4TPV). The secondary-structure cores are highly superimposable across the three domains, whereas the largest RMSD values localize to surface-exposed loops. These loops correspond with the regions of highest predicted antigenicity, indicating that isoform- and domain-specific immunoreactivity is driven by surface divergence.

**Fig. 3.**
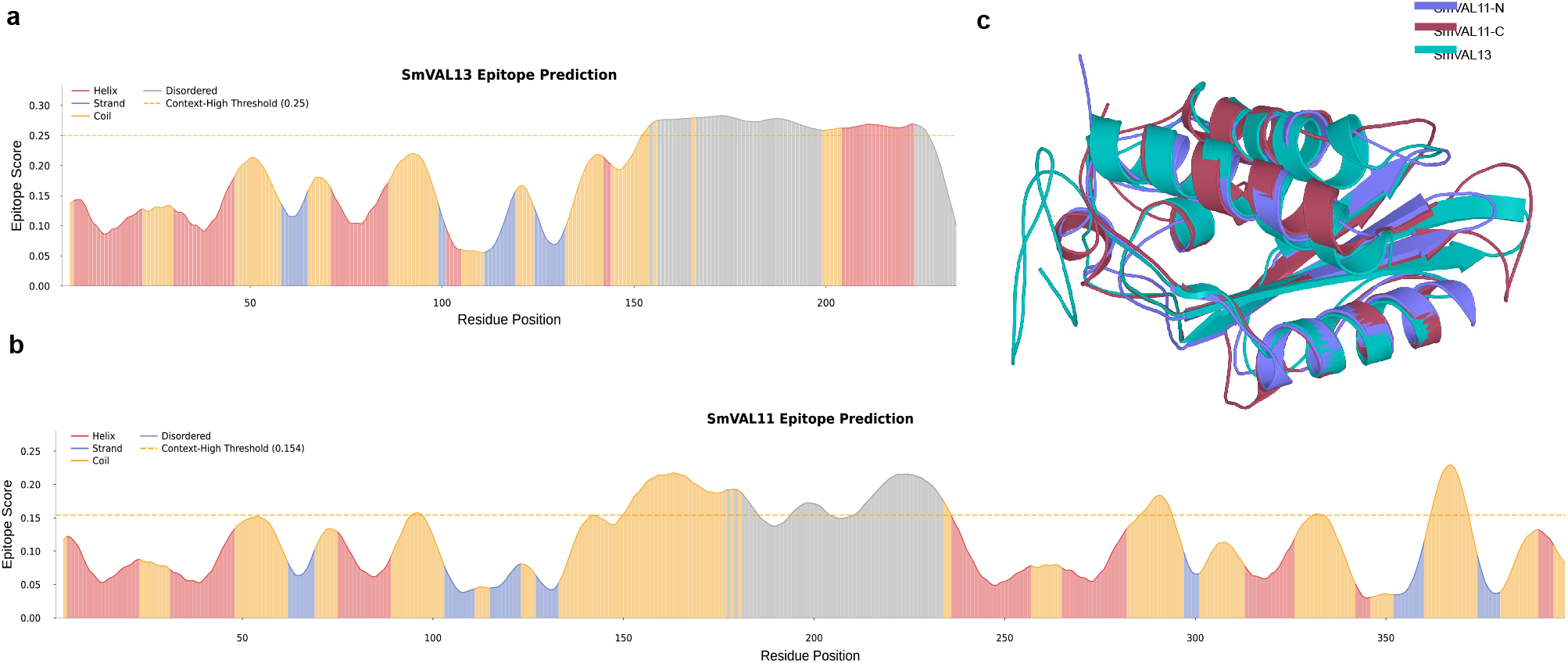
Prediction of antigenicity and three-dimensional fold of SmVAL13 and SmVAL11. (a) BepiPRED antigenicity prediction combined with DISOPRED and PSIPRED of SmVAL13. (b) BepiPRED antigenicity prediction combined with DISOPRED and PSIPRED of SmVAL11. (c) Superposition of the three homology models: SmVAL11-N (purple), SmVAL11-C (dark wine) and SmVAL13 (cyan). The low complexity and intrinsically disordered regions are in agreement with that predicted to be highly antigenic.

### Antibody production and characterization

Following the prediction of absence of cross reaction of anti-SmVAL antibodies, we raised isoform-specific polyclonal antibodies (pAbs). Polyclonals were chosen for their multi-epitope recognition, which well matches to the flexible surface epitopes identified and their low cost. We outsourced the pAb production and 1 rabbit per protein was immunized and one received adjuvant as a control. Intermediate bleeds were tested, and a positive response was clear for both antigens, starting from the first bleed at 14 days post injection. At the end of the 3 injections only one rabbit per protein was finally sacrificed. The pAb anti-SmVAL11 and anti-SmVAL13 were purified from rabbit serum by affinity chromatography on protein A column, concentrated to 3 mg/ml and stored at −20 °C in aliquots containing 20% glycerol. The purified pAb were then tested by western blot (Fig. 1a and 2a) on the purified recombinant proteins and on red blood cells from non-infected blood (Fig. S1). pAb anti-SmVAL13 and pAb anti-SmVAL11 were tested against recombinant SmVAL13 and SmVAL11 to verify sensitivity and cross-recognition. Each pAb detected its cognate antigen down to a 1:10,000 dilution; moreover, they do not cross-react with each other up even at 1:500 dilution (Fig. S1), confirming the isoform specificity predicted from the epitope analysis. Critically, none of the two pAb recognized human proteins in non-infected blood samples up to 1:500 dilution (Fig. S1), establishing a clean background against the sample matrix.

### Antibody-antigen complex characterization by SAXS and BLI

To understand the architecture and stoichiometry of the pAb–antigen complexes, we combined SEC-SAXS and biolayer interferometry (BLI). For SEC-SAXS we prepared three pAb:SmVAL ratios (1:0.5, 1:1, 1:2), with the pAb alone as control (Fig. 4a, 4b, burgundy curve). The free antibodies showed the characteristic IgG profile, with a main P_r_ distribution near 3 nm and a shoulder at ∼7.5 nm (Fig. 4c, 4d, burgundy curve). Both pAbs gave similar parameters (Table 3). When the cognate complexes were analyzed, a larger, diffuse peak appeared at higher sizes, growing in intensity with antigen ratio. Although it appeared as a double population, EFA resolved it as a single, broad species. In both complexes (pAb:SmVAL13 and pAb:SmVAL11) this large population reflects simultaneous engagement of multiple epitopes by the polyclonal mixture, which cross-links antigen and antibody into large heterogeneous assemblies. For the pAb:SmVAL11 complex, an additional smaller peak matching with unbound protein appeared and increased with SmVAL11 excess.

**Fig. 4.**
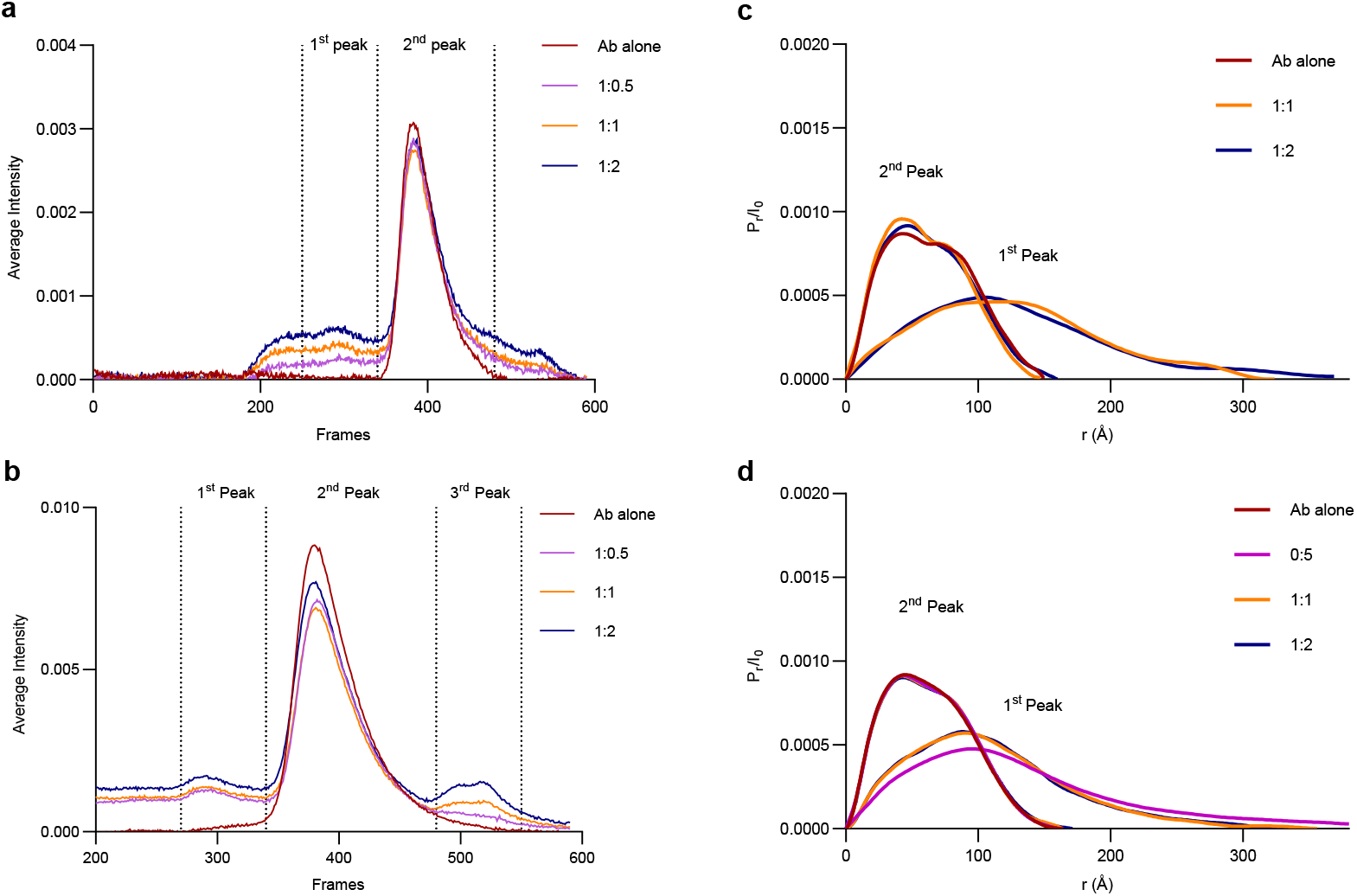
SEC-SAXS analysis of the complexes between polyclonal Ab and SmVALs. (a) Superposed SEC-SAXS chromatograms SmVAL13 in complex with its cognate Ab: Ab alone (red); Ab:SmVAL13 (1:0.5) (violet); Ab:SmVAL13 (1:1) (light orange); Ab:SmVAL13 (1:2) (blue). (b) Superposed SEC-SAXS chromatograms of Ab-anti SmVAL11 alone (red) and in complex with SmVAL11 in different ratios: 1:0.5 (violet), 1:1 (light orange) and 1:2 (blue). (c) Pair distance distribution graph of the eluted peaks from (a) after EFA deconvolution. (d) Pair distance distribution graph of the eluted peaks from (b) after deconvolution.

**Table 3.** Summary of SEC-SAXS analysis of pAb-antiSmVAL13, pAb-antiSmVAL11 and their complexes with cognate proteins. The chromatographic peaks were deconvoluted with EFA and the scattering parameters were calculated with ATSAS (Rg and Dmax) and Bayesian inference (MW).

| Sample (peak) | Ratio | $R_g$ (nm) | $D_{max}$ (nm) | MW (kDa) |
| --- | --- | --- | --- | --- |
| pAb anti-SmVAL13 | - | $5.05 \pm 0.01$ | 16.2 | 146.8 |
| pAb:SmVAL13<br>(1<sup>st</sup> peak) | **1:1** | $10.1 \pm 0.24$ | 32.4 | 646.8 |
| | **1:2** | $10.5 \pm 0.34$ | 36.9 | 646.8 |
| pAb:SmVAL13<br>(2<sup>nd</sup> peak) | **1:0.5** | $4.77 \pm 0.02$ | 14.3 | 124.5 |
| | **1:1** | $4.74 \pm 0.02$ | 14.7 | 129.1 |
| | **1:2** | $4.97 \pm 0.02$ | 16.0 | 138.2 |
| pAb anti-SmVAL11 | - | $4.92 \pm 0.02$ | 16.4 | 146.8 |
| pAb:SmVAL11<br>(1<sup>st</sup> peak) | **1:0.5** | $11.1 \pm 0.21$ | 42.7 | 518.4 |
| | **1:1** | $8.87 \pm 0.25$ | 35.6 | 479.1 |
| | **1:2** | $8.81 \pm 0.13$ | 33.2 | 433.6 |
| pAb:SmVAL11<br>(2<sup>nd</sup> peak) | **1:0.5** | $4.94 \pm 0.01$ | 16.1 | 140.9 |
| | **1:1** | $4.98 \pm 0.01$ | 16.4 | 146.8 |
| | **1:2** | $5.01 \pm 0.01$ | 17.2 | 146.8 |

The binding of each recognition element to its cognate partner was characterized by BLI. The two polyclonal antibodies engaged their antigens with incomplete and slow dissociation kinetics. The anti-SmVAL13 antibody displayed k_off_ = 2.9 × 10^-4^ ± 0.5 × 10^-4^ s^-1^, whereas the anti-SmVAL11 antibody dissociated about five-fold faster and with a broader spread (k_off_ = 1.6 × 10^-3^ ± 0.6 × 10^-3^ s^-1^), as shown in Fig. S5. This faster and more dispersed apparent off-rate is consistent with the heterogeneous nature of SmVAL11, in which each population dissociates independently, but concomitantly. For both antibodies the corresponding apparent equilibrium dissociation constants lay in the nanomolar range (Table S1). Overall, the kinetic mechanism is compatible with an avidity-driven recognition, with a fast k_on_ and a slow and incomplete k_off_.

### Molecularly Imprinted Polymers binding to SmVAL13

To overcome the costs for raising Ab, whether monoclonal or polyclonal, we decided to explore a synthetic alternative offered by the Molecularly Imprinted Polymers (MIP). Given the proneness of SmVAL11 to auto-cleave into two separate domains, we decided, for a proof of concept, to target first SmVAL13, which was conjugated *via* the N-terminus to the aldehyde groups on the magnetic nanoparticles (MNP) surface. Subsequently, SmVAL13-specific nanoMIPs were prepared by polymerizing acrylamide-based functional monomers onto SmVAL13 functionalized MNPs. Once released from the MNP surface, the nanoMIP was lyophilized. The MNPs were recycled for two further nanoMIP synthesis cycles (60 min per cycle). The average nanoMIP yield per cycle was determined to be 8.8 mg ± 0.9 mg (Fig. 5a). For multi angle dynamic light scattering (MADLS) characterization, 1 mg of nanoMIP was resuspended in 1 mL of PBS, followed by a 1:10 dilution in ultrapure water. Particle diameter was determined to be 176 nm ± 23 nm with a polydispersity index of 1.0 indicating a monodisperse ideal distribution of nanoMIP particles (Fig. 5b). Particle concentration at a diameter of 176 nm was determined to be around 3 x10^10^ particles/ml as shown in Fig. 5b.

**Fig. 5.**
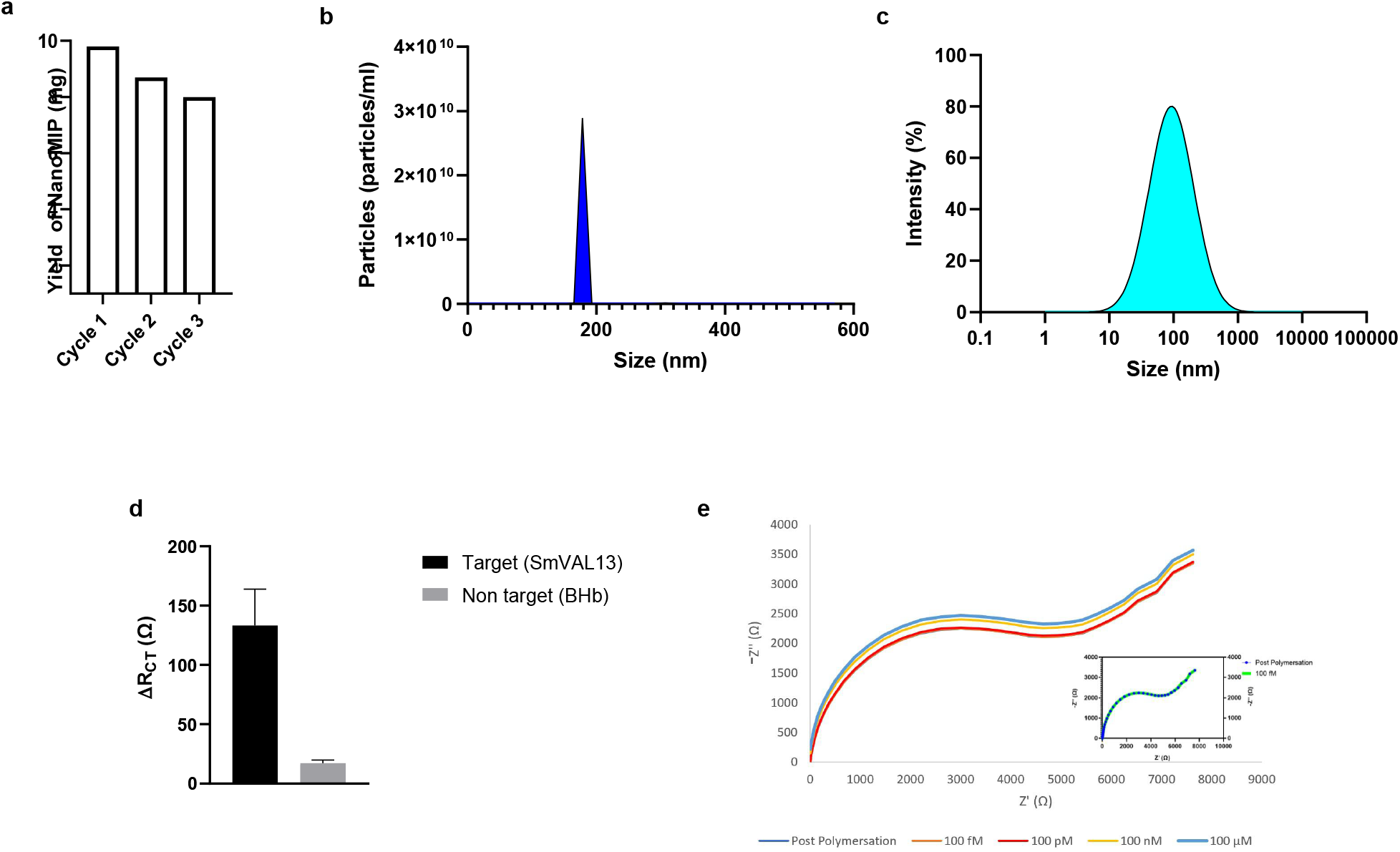
Production, characterization of nanoMIPs and testing against SmVAL13. (a) Lyophilized yield over 3 cycles of SmVAL13 nanoMIP synthesis; (b) MADLS determination of nanoMIP particle size; (c) MADLS determination nanoMIP particle concentration (d) EIS protein selectivity study using SmVAL13 vs non-target (BHb) at 1nM of protein binding; (e) Nyquist plots obtained following polymerization and after exposure to increasing target concentrations (100 fM–100 μM). The inset highlights the overlap between the post-polymerization baseline response (green line) and the 100 fM response (blue line with markers), indicating minimal impedance change at the lowest concentration tested.

NanoMIPs (5 µg) suspended in 50 µL were electrochemically entrapped on screen-printed gold electrodes using cyclic voltammetry and the resulting modified surface was used as a biosensor to determine selective protein attachment from standard solutions of target SmVAL13 and non-target bovine hemoglobin (BHb), as shown in Fig. 5d. Electrochemical impedance spectroscopy (EIS) was used to determine changes in the charge transfer resistance (ΔR_CT_) upon target and non-target binding at 1 nM. Fig. 5d shows the difference in signal for target (SmVAL13) *vs* non-target (BHb) demonstrating that the nanoMIP-based sensor was selective for SmVAL13 at a ratio of 10:1. Additionally, the SmVAL13 sensor was used to determine a calibration range for target binding and subsequently to determine an equilibrium dissociation constant (K_D_) for SmVAL13 (Fig. S4). The sensor was able to determine SmVAL13 over a wide concentration range (100 fM to 100 µM, Fig. 5e). LOD and LOQ were determined to be 100 fM and 1 pM respectively. The dissociation constant (K_D_) was determined to be in the low nanomolar range (K_D_= 87.7 ± 0.037 nM) averaged across the three runs using the Hill–Langmuir adsorption isotherm model using data from Fig. S4. In this approach, K_D_ is defined as the concentration of target protein at half maximal binding (B_max_/2), with non-specific binding minimized through washing steps designed to remove loosely or non-specifically associated material. The analysis assumes a 1:1 binding interaction between the target protein and the binding sites.

In BLI, the SmVAL13-imprinted nanoMIP recognized its target with a dissociation rate constant k_off_ = 8.9 × 10^-3^ ± 1.6 × 10^-3^ s^-1^ (Figure S5), roughly one order of magnitude faster than either antibody, moreover the dissociation is almost complete. The distinguishing feature of the nanoMIP relative to the pAb is therefore kinetic rather than in overall affinity (K_D_= 97 ± 35 nM): it combines nanomolar-range recognition with a substantially faster off-rate (Table S1). Such a reversible non-avid interaction is a favorable property for a reusable biosensor, as it facilitates surface regeneration between measurements.

## Discussion

SmVAL13 and SmVAL11 are circulating SCP/TAPS proteins, presenting biophysical and structural properties that make them suitable biomarkers for detecting schistosomiasis infection in human blood sample. In particular, they can be expressed in high yield in a bacterial heterologous system, they are monodispersed, thermally stable and show high antigenicity.

From a structural standpoint, SmVAL13 and SmVAL11 exhibit distinct domain organizations that might reflect their roles in different parasite stages. SmVAL13 is characterized by one single SCP/TAPS domain, as confirmed by circular dichroism spectroscopy. Its thermal melting of 57 °C, suggests good stability and resistance to thermal stress. The overall flexibility, inferred from the elongated conformation observed in SAXS and the larger particle volume in DLS, suggests a dynamic structure in solution, possibly contributing to its antigenic properties. This conformational flexibility, particularly in the C-terminal region, coincides with its predicted antigenic sites, indicating a potential use as biomarker for immune recognition.

SmVAL11 contains two duplicated domains connected by an *in vitro* auto-cleavable linker. Despite this modular organization, the protein behaves as a single unit, as confirmed by DLS and SAXS. This supports the hypothesis that the two individual SCP/TAPS domains could interact in solution, independently whether full length or cleaved.

Polyclonal antibody production revealed a very good immunogenicity of both proteins. Furthermore, they do not cross-react with each other and do not recognize human proteins found in blood. The pAb:SmVALs complexes exhibited high flexibility, suggesting multiple recognition sites on the antigen and/or conformational states. A low-resolution *ab-initio* model built by DAMMIF from the SAXS data of a 1:1 stoichiometry is proposed in Fig. S2, where polyclonal Ab seems to bind the initial part of the disordered region of SmVAL13, whose *ab-initio* SAXS model was docked into the complex. This will be used as a starting point to dissect SmVAL13 sequence for epitope mapping.

NanoMIPs exhibited high sensitivity toward SmVAL13, with binding affinities similar to the cognate polyclonal raised, while maintaining strong target selectivity. NanoMIP demonstrates affinities akin to monoclonal antibodies. Its compatibility with and facile integration onto disposable electrode platforms further supports their utility in point-of-care and in-field diagnostic applications. In addition, nanoMIPs can be produced at substantially lower cost than their antibody counterparts (approximately £1230 for Ab production versus £60 for nanoMIP, as summarized in Table S1), highlighting a significant economic advantage. Notably, these materials retain functional stability following prolonged storage at room temperature, remaining active after several months without degradation. Taken together, the combination of superior binding performance, speed of production, low production cost, and robust stability under non-refrigerated conditions positions nanoMIPs as highly promising candidates for widespread biosensing of SmVAL13 as a disease biomarker, particularly in settings where financial resources and cold-chain infrastructure are limited.

In conclusion, the biophysical characterization of SmVAL13 and SmVAL11 alone and in complex with their cognate polyclonal Ab and nanoMIP not only highlights their roles as potential biomarkers for *S. mansoni* infection, but also provides a foundation for the rational design of a diagnostic kit. Future work will focus on refining the structural models and exploring their diagnostic efficacy in clinical samples, with the ultimate goal of providing a rapid, cost-effective, and non-invasive diagnostic solution for schistosomiasis detection.

## Methods

### Materials

Unless specified, all chemicals and buffer powders were purchased from Sigma-Aldrich-Merck and were of analytical grade. FeCl_3_·6H_2_O, sodium acetate (NaOAc), glutaraldehyde, ethylene glycol, N-hydroxymethylacrylamide (NHMA), N,N′-methylenebisacrylamide (MBAm), sodium dodecyl sulfate (SDS), ammonium persulfate (APS), tetramethylethylenediamine (TEMED), potassium persulfate (KPS), and sodium nitrate (NaNO_3_), dithiothreitol (DTT) and methylhydroquinone (MHQ) were used as received without further purification. Ultrapure water (18.2 MΩ cm) was obtained from a Milli-Q purification system. Phosphate-buffered saline (PBS) consisted of 10 mM phosphate, 137 mM NaCl and 2.7 mM KCl, adjusted to pH 7.4. Metrohm Dropsense disposable screen-printed electrodes (Au-BT 220) comprising a gold working electrode (0.4 cm diameter), a gold counter electrode and silver reference electrode were purchased from Metrohm (Runcorn, Cheshire, UK).

### Protein expression and purification

SmVAL11 and SmVAL13 synthetic genes (GeneArt, DB) were cloned into pTXB1 (New England Biolabs) and transformed into E. coli BL21(DE3) Gold (ThermoFisher Scientific) for expression. A 50 mL LB culture with 50 µg/mL ampicillin was grown overnight at 37 °C (shaking at 130 rpm), diluted 1:100 into 1 L LB-ampicillin, grown to Abs600 = 0.6–0.8, and induced with 0.2 mM IPTG (Sigma-Aldrich) at 20°C for 16 h (shaking at 130 rpm). Cells were harvested (4,000 × g, 20 min, 4 °C), resuspended in lysis buffer [50 mM Tris-HCl, pH 8.5, 500 mM NaCl, 1 mM EDTA, 5 U DNase, EDTA-free protease inhibitor (Roche)], and lysed by sonication (12 × 60 s, 50% amplitude, on ice). The clarified lysate (15,000 × g, 60 min, 4°C) was loaded onto a chitin resin column (IMPACT Kit, New England Biolabs) equilibrated with lysis buffer, washed with 10 column volumes each of wash buffer 1 (50 mM Tris-HCl, pH 8.5, 500 mM NaCl, 1 mM EDTA) and wash buffer 2 (25 mM MES, pH 6.8, 125 mM NaCl, 1 mM EDTA), and cleaved with buffer 3 (25 mM MES, pH 6.8, 125 mM NaCl, 1 mM EDTA, 60 mM DTT) for 16 h at 4 °C. The eluted protein was dialyzed 1:200 against storage buffer (25 mM MES, pH 6.8, 125 mM NaCl, 1 mM EDTA) using a 3 kDa MWCO membrane, concentrated (Amicon Ultra-15, 10 kDa MWCO), and stored at −80 °C. Purity was assessed by 15 % SDS-PAGE with Coomassie staining, and identity was confirmed by Western blotting using anti-SmVAL13 and anti-SmVAL11 pAb.

### BepiPred and Homology modeling

Linear epitope prediction was performed using BepiPred 3.0 [Clifford et al., 2022] using a high context threshold (75% signal threshold). Secondary structure prediction was carried out using PSIPRED 4.0 [Buchan & Jones, 2019], while intrinsic disorder was assessed with DISOPRED3 [Jones & Cozzetto, 2015]. All three predictions were performed on the full-length sequences of SmVAL13 and SmVAL11. Results were combined and visualized as a single composite figure, mapping epitope scores, predicted secondary structure elements (α-helix, β-sheet, random coil) and disorder probability along the sequence, to correlate antigenicity with structural and flexibility features of each protein (Figure 3a, 4b).

Three-dimensional models of SmVAL11-N, SmVAL11-C and SmVAL13 were built *via* the MOE homology modeling program (Molecular Operating Environment, Chemical Computing Group, Montreal, QC, Canada). SmVAL4, SmVAL11-N, SmVAL11-C and SmVAL13 were aligned using Clustal Omega [Madeira et al., 2024] as shown in Supplementary fig. 2. The PDB Search program of MOE identified a set of homologous templates (PDB IDs: 4AIW.A – human GAPR1, 5ETE.A, 1CFE.A, 5V50.A - *Moniliophthora perniciosa* MpPR-1i, 4TPV.A - *Ancylostoma caninum* Hookworm Platelet Inhibitor), belonging to precomputed domain-based PDB families, defined using PFAM as a guide. SmVALs and the pre-aligned templates were kept as two separate blocks, with the internal alignment of each block fixed, and the two blocks were then reciprocally aligned. SmVAL11-N and SmVAL11-C three-dimensional models were built using 4AIW.A as template. SmVAL13 was modeled using 4AIW.A as main template and the C-terminal residues of 4TPV.A as secondary template, allowing us to extend the three-dimensional model of the C-terminal disordered tail.

### Dynamic light scattering (DLS)

SmVAL13 and SmVAL11 were diluted to 1 mg/mL in storage buffer (25 mM MES, pH 6.8, 125 mM NaCl, 1 mM EDTA) and filtered (0.22 µm). Measurements were performed using a Malvern Zetasizer Pro (Malvern Panalytical, Malvern, UK) at 25 °C in a low-volume quartz cuvette. Three runs of 10 acquisitions (120s equilibration, 10 s each) were collected per sample, with the laser wavelength of 633 nm and detector in backscattering position. Hydrodynamic radius and polydispersity index were calculated using Zetasizer software (v8.0).

### Circular Dichroism (CD)

CD spectroscopy was performed using a Chirascan spectrometer (Applied Photophysics) equipped with a Peltier temperature control unit. SmVAL13 and SmVAL11 were analyzed at 10 µM in 2.5 mM MES pH 6.8, 12,5 mM NaCl, and filtered (0.22 µm). Samples were loaded into a 1 mm path length quartz cuvette (Hellma Analytics, Müllheim, Germany). Far-UV spectra (185–260 nm) were collected at 20°C with a 0.5 nm step size, 0.5 s integration time, 1 nm bandwidth and three accumulations per spectrum. Buffer baselines were recorded under identical conditions and subtracted. Thermal stability was assessed by monitoring ellipticity at 222 nm from 20°C to 80°C (1°C/min ramp, 1°C steps, 3 spectra/temperature). Buffer subtracted and averaged spectra were smoothed using a Savitzky-Golay filter (5-point window).

### Small Angle X-Ray Scattering

SEC-SAXS experiments were conducted at the SWING beamline, SOLEIL synchrotron (Saint-Aubin, France), batch mode SAXS was performed at ESRF BioSAXS Bm29 beamline (Grenoble, France). SmVAL13 was concentrated to 1.2, 2.4, 3.6, and 7.3 mg/mL for experiments in batch mode, whereas in SEC-SAXS, in complex with pAb-anti SmVAL13 the concentrations were 10 µM (0.3 mg/mL), 20 µM (0.6 mg/mL) and 40 µM (1.2 mg/mL). pAb-anti SmVAL13 concentration was kept at 20 µM (2.8 mg/mL). SmVAL11 was concentrated to 11 mg/mL for the batch analysis. In complex with pAb-anti SmVAL11 the concentrations of SmVAL11 were 12,5 µM (0.55 mg/mL), 25 µM (1.1 mg/mL) and 50 µM (2.2 mg/mL). pAb anti-SmVAL11 concentration was kept at 25 µM (3.5 mg/mL). Matching buffer was prepared by dialysing storage buffer against protein-free buffer to minimize scattering mismatch. Samples and buffers were loaded into a 1.5 mm diameter quartz capillary in a vacuum chamber, maintained at 20 °C. Data were collected using a monochromatic X-ray beam [λ = 0.76 Å (SOLEIL) and 0.99 Å (ESRF)]. Buffer scattering was measured before and after each sample and averaged for subtraction. Data were inspected for radiation damage by comparing consecutive frames; damaged frames were excluded. Raw data were reduced using FOXTROT (v3.5). Buffer subtraction and Evolution Factor Analysis and data refinement was done using RAW [Hopkins, 2024]. Guinier analysis was performed to estimate radius of gyration (R_g_). Pair-distance distribution functions Pr were calculated using GNOM (ATSAS v3.0.4, [Manalastas-Cantos et al., 2021], and low-resolution molecular envelopes were generated with DAMMIF (14 independent runs, averaged with DAMAVER, from the ATSAS suite). Deconvolution of SEC-SAXS peaks was performed with EFA software [Hopkins JB. 2024], according to the online documentation.

### Mass spectrometry analysis

Electrophoretic bands corresponding to SmVAL11 were cut from the gel and submitted to tryptic proteolysis. Briefly, the selected bands were washed several times with an aqueous solution of 50 mM ammonium bicarbonate, destained with 50% acetonitrile:bicarbonate solution and dehydrated at the end with pure acetonitrile. Then they were reduced with 10 mM dithiothreitol (DTT) and alkylated with 55 mM iodic-acetamide (IAA). The proteolysis was carried out with 100 ng of trypsin in 25 mM ammonium bicarbonate at 37 °C overnight. MALDI-ToF MS analyses were performed with an AutoFlex II instrument (Bruker Daltonics, Bremen, Germany), equipped with a 337 nm nitrogen laser and operating in reflector positive mode.

### Production of polyclonal antibodies against SmVAL11 and SmVAL13

10 mg of pure SmVAL11 and SmVAL13 were sent to Cambridge Research Biochemicals (CRB, UK). 1 rabbit was immunized against each recombinant protein and 1 received a control solution with the adjuvant only. 3 successive immunizations were performed at 14 days interval each and test bleeds were taken one week after each immunization. For both SmVAL11 and SmVAL13 the bleed of only one of the two rabbits was effective in recognizing the recombinant protein. The harvest bleed was collected one week after the last immunization only from the positive rabbit. All the positive bleeds were pooled together, and the Abs were purified by affinity chromatography on protein A column (Cityva), according to manufacturer instructions. The purified Abs were then concentrated by ultrafiltration with a 30 kDa MWCO, added with 20% glycerol, aliquoted and stored at −20 °C until use.

### Western blot analysis

Western blot analysis after SDS-PAGE was performed on polyvinylidene difluoride (PVDF) membranes (BioRad), activated by 2 min soaking in methanol. Towbin buffer (25 mM Tris, 192 mM glycine, 20% ethanol, 0.1% SDS) was used for transferring the protein bands into the PVDF membrane in a wet transfer apparatus (Hoefer). The transfer was allowed for 1:30 h at 4 °C under a constant current intensity of 400 mA.The membrane was incubated for 1:30h at room temperature in TBS-T (20 mM Tris/HCl pH 7.6, 137 mM NaCl, 0.1% Tween-20) supplemented with 5% dried milk, to avoid any unspecific binding. Three consecutive washes of 5 min each with TBS-T preceded and followed the incubation with the primary Ab (1:10000 or 1:500), overnight at 4 °C. The Ab anti-SmVALs was revealed by using a HRP-conjugated mouse anti-Rabbit Ab (Sigma), diluted 1:10000 for 1.30 h. The HRP was revealed by chemiluminescence with Immobilon Crescendo western blot kit (Millipore); images were collected with ChemiDoc (BioRad).

### Biolayer interferometry (BLI)

Interaction experiments were performed on an OctetRED96 system (Pall, ForteBIO) using amine-reactive second-generation (AR2G) biosensors (Sartorius). Sensors were hydrated in water for 10 min, then carboxylic groups were activated for 10 min with 0.1 M N-hydroxysulfosuccinimide (NHS) and 0.2 M 1-ethyl-3-(3-dimethylaminopropylcarbodiimide hydrochloride (EDC). SmVAL13 (100 nM), pAb (100 nM), or nanoMIP (20 or 40 µg/ml) were covalently coupled to the sensor surface via their primary amine groups in 10 mM sodium acetate pH 5.4 for 10 min. Residual activated groups were blocked with 1 M ethanolamine pH 8.5 for 10 min. The sensors were equilibrated in 50 mM MES pH 6.8 for 60 s, then dipped into solutions containing the binding partners at increasing concentrations of 160, 250, 320, 500, and 700 nM (with an additional 100 nM point for the nanoMIP surfaces). Association and dissociation profiles were reference-subtracted and processed with the manufacturer’s data analysis software. Binding curves were fitted globally, across the association and dissociation phases of the full concentration series, to a 2:1 heterogeneous-ligand model, which returned two sets of rate constants (k_on1_/k_off1_ and k_on2_/k_off2_) and the corresponding equilibrium dissociation constants K_D1_ and K_D2_ (Supplementary Table S1). For the immobilized antibody the first component was systematically below the instrument resolution limit (K_D1_ < 1 pM). For the immobilized nanoMIP the two components were not kinetically resolved (k_on1_ ≈ k_on2_), indicating a more plausible 1:1 binding mode, nevertheless for better comparison we decided to keep the same binding model all throughout. To ensure that only well-determined cycles contributed to the averaged parameters, each individual fit was retained solely when the fit quality was R² ≥ 0.95 and the second-component dissociation rate exceeded 10 ^-5^ s^-1^ (a genuinely resolved, dissociating second component); cycles failing either criterion were discarded. Retained kinetic parameters (K_D2_, k_off2_) are reported as mean ± SD across the retained concentrations for each sensor. Independently, a steady-state equilibrium dissociation constant (K_D_) was derived for each sensor from a Langmuir fit of the equilibrium response as a function of analyte concentration; this steady-state K_D_ is a distinct quantity from the kinetic K_D2_ and the two are reported separately. Steady-state fits with R² < 0.90 were excluded from group-level averaging.

### Molecularly Imprinted Polymer production and quality control

#### Magnetic nanoparticle synthesis

Aldehyde-functionalized magnetic nanoparticles (MNPs) were synthesized using a previously reported solvothermal microwave-assisted method (Stephen et al., 2025; Sullivan, Stockburn, et al., 2021). FeCl_3_·6H_2_O (0.5 g) and NaOAc (1.8 g) were dissolved in ethylene glycol (15 mL) in a 30 mL microwave reaction vial (G30, Anton Paar) under magnetic stirring. To impart aldehyde-functionalisation (MNP@CHO), glutaraldehyde (3.5 mL) was added and the reaction mixture stirred for a further 5 min at room temperature. The stirrer bar was removed and the sealed vial transferred to a Monowave 200 microwave reactor (Anton Paar). Samples were heated to 200 °C at a ramp rate of 18 °C min-1 (10 min ramp time) and maintained at 200 °C for 20 min under autogenous pressure (∼9 bar).

Following synthesis, reaction mixtures were cooled for 10 min at room temperature to 70 °C. Nanoparticles were magnetically isolated using a neodymium magnet and washed five times with Ultrapure water (18.2 MΩ cm) followed by two washes with absolute ethanol. Purified particles were resuspended in Ultrapure water at a final concentration of 300 mg in 30 mL and stored at 4 °C until further use.

#### Protein functionalization of magnetic nanoparticles

Aldehyde-functionalized nanoparticles (MNP@CHO; 1 mL suspension corresponding to 10 mg particles) were transferred to 1.5 mL microcentrifuge tubes and magnetically separated for 10 min using a neodymium magnet. The supernatant was discarded and replaced with 1 mL of recombinant SmVAL13 solution (1 mg mL-1) prepared in MES storage buffer (25 mM MES, 125 mM NaCl, 1 mM EDTA, pH 6.8). Followed by vigorous vortex mixing to ensure homogeneous suspension. The reaction mixture was incubated for 30 min at room temperature (22 °C) to allow conjugation of SmVAL13 to the aldehyde-functionalized nanoparticle surface.

Following incubation, particles were magnetically separated and washed three times with fresh PBS buffer three times. Protein concentration measurements were performed spectrophotometrically using a BioDrop μLITE UV–visible spectrophotometer. Protein immobilization efficiency was determined by comparing the initial protein concentration with the residual protein concentration remaining in the supernatant following conjugation resulting in 0.6 mg of SmVAL13 being up taken per 10 mg of MNP@CHO. SmVAL13-functionalized nanoparticles (MNP@CHO@SmVAL13) were stored in aqueous suspension at 4 °C until further use.

#### NanoMIP synthesis

Molecularly imprinted polymer nanoparticles were synthesized directly upon the surface of MNP@SmVAL13 templates by free-radical polymerization. Magnetic nanoparticles were dispersed in PBS (906 µL, pH 7.4) by sequential sonication, vigorous shaking and vortex mixing before transfer to a 15 mL Falcon tube. Samples were mixed at 400 rpm at room temperature using a thermomixer and degassed under nitrogen for 15 min. Following degassing, NHMA (54 µL), SDS (0.4 mg) and MBAm (6 mg) were added to the reaction mixture. Polymerization was initiated immediately by adding 20 µL of an aqueous initiator solution containing 10% (v/v) TEMED and 5% (w/v) APS. A nitrogen headspace was introduced and the reaction vessel sealed with the cap before incubation at 400 rpm for 15 min at room temperature.

Polymerization was quenched by addition of MHQ (1 mL, 10 mM) followed by vigorous vortexing. Reaction mixtures were then washed three times with fresh PBS to remove residual monomer, surfactant and quencher. Following magnetic separation, MNP@CHO@SmVAL13∼nanoMIP composites were resuspended in ultrapure water (600 µL) and subjected to ultrasonication (VWR ultrasonic bath, 600 W, 45 kHz) at 37 °C for 5 min to release the non-covalently surface-bound nanoMIPs from the magnetic templates. Magnetic particles (MNP@CHO@SmVAL13) were removed using a neodymium magnet and re-used for a further two cycles of nanoMIP synthesis. The supernatant containing released nanoMIPs was collected and stored at 4 °C until further use.

#### NanoMIP lyophilization and yield determination

NanoMIP suspensions were flash-frozen in liquid nitrogen prior to lyophilization using a CHRIST Alpha 2–4 LDplus freeze dryer. Microcentrifuge tube caps were left open and covered with perforated Parafilm® to permit solvent sublimation during freeze drying. Samples were lyophilized at −90 °C under reduced pressure (0.011 mbar) for a minimum of 16 h until a fine off-white powder was obtained. NanoMIP yield (8.8 mg ± 0.9 mg) was determined gravimetrically by mass balance following lyophilization. Lyophilized nanoMIPs were stored at 4 °C until further use.

#### Dynamic light scattering and particle concentration analysis

Hydrodynamic diameter, dispersity and particle concentration measurements were performed using a Zetasizer Pro Red Label instrument (Malvern Panalytical, Malvern, UK) equipped for multi-angle dynamic light scattering (MADLS). Samples were analysed in aqueous suspension at a 1:10 dilution in ultrapure water at 25 °C according to the manufacturer’s instructions.

#### Binding analysis and dissociation constant determination

Binding analysis and dissociation constant calculations were performed using GraphPad Prism (GraphPad Software, v10). Binding data were fitted using a one-site specific binding model based on Langmuir adsorption kinetics. Non-specific binding contributions were minimized through sequential washing steps prior to each measurement and were therefore not incorporated into the final fitting model.

#### Electrochemical deposition of nanoMIP-containing films on screen-printed electrodes

All electrochemical experiments were performed using a PGSTAT204 potentiostat (Metrohm Autolab) controlled by NOVA v2.1.4 software. Gold screen-printed electrodes (BT-Au SPEs; Metrohm), incorporating a silver/silver chloride (Ag/AgCl) reference electrode, were used as received. The geometric working electrode diameter was 4 mm. NanoMIP-containing polymer layers were fabricated via cyclic voltammetry (CV), adapted from previously reported methods (Reddy et al., 2024). Briefly, a 50 μL polymerization solution in PBS containing nanoMIPs (0.1 mg), NHMA (1.33 M), MBAm (41.5 mM), NaNO₃ (0.29 M), and KPS (48.15 mM) was drop-cast onto the electrode surface. Electropolymerization was performed by cycling the potential between −0.2 V and −1.4 V (vs. Ag/AgCl) for 7 cycles at a scan rate of 50 mV s⁻¹ (∼10 min total) at room temperature (22 ± 2 °C). This process yielded E-layers with entrapped nanoMIP islands (E-NMI). Control E-layers were prepared under identical conditions in the absence of nanoMIPs. No degassing or surface blocking steps were required prior to electropolymerization. Following polymerisation electrodes and the entrapped nanoMIP layer electrochemically conditioned in PBS (50 μL) sweeping the potential between −0.5 V and +1.5 V (vs. Ag/AgCl) for 5 cycles at a scan rate of 175 mV s⁻¹ (∼5 min total) at room temperature (22 ± 2 °C).

#### Protein rebinding and selectivity assays

NanoMIP-containing E-layers (E-NMI) and control electrodes were incubated with solutions of the target protein SmVAL13 over a concentration range of 100 fM to 100 μM. All protein solutions were prepared in PBS. Each incubation was conducted for 5 min at room temperature (22 ± 2 °C), followed by rinsing with PBS to remove unbound protein prior to electrochemical analysis. Selectivity was assessed by performing identical incubations using haemoglobin as a non-target protein. All binding and selectivity experiments were performed in triplicate (n = 3).

#### Electrochemical impedance spectroscopy (EIS)

EIS measurements were carried out using a solution consisting of 5 mM potassium ferricyanide in PBS supplemented with 0.5 M KCl as supporting electrolyte. The measurements were conducted at an applied potential of 0.1 V (± 0.01 V) with a sinusoidal perturbation amplitude of 10 mV over a frequency range of 0.1 Hz to 100 kHz. Impedance spectra were acquired across 10 frequency decades. Data were analyzed using a Randles equivalent circuit model implemented in the FRA32 module (Metrohm Autolab). The charge-transfer resistance (Rct) was extracted as the primary analytical parameter (see Supplementary Fig. S4).

#### Analytical performance and statistics

The limit of detection (LOD) and limit of quantification (LOQ) for SmVAL13 were determined from the calibration curve generated by plotting the change in charge-transfer resistance (ΔR_CT_) against the logarithm of analyte concentration. A sigmoidal response was observed, with increasing ΔR_CT_ values corresponding to increasing SmVAL13 concentrations. The LOD was defined as the lowest concentration that produced a measurable response significantly greater than the baseline signal following post-polymerisation. Based on the calibration curve, a detectable change in ΔR_CT_ was first observed at 100 fM, which was therefore assigned as the sensor’s LOD. The LOQ was determined as the lowest concentration that could be quantified with acceptable accuracy and reproducibility, corresponding to the beginning of the linear response region of the calibration curve. From the fitted response, this occurred at 1 pM and was therefore assigned as the LOQ. LOD and LOQ values were estimated using the standard analytical expressions: LOD = 3σ/*m* and LOQ = 10σ/*m* where σ is the standard deviation of the blank or baseline response and *m* is the slope of the linear region of the calibration curve. The obtained results demonstrate the high sensitivity of the EIS sensor platform toward SmVAL13 detection at sub-picomolar concentrations. All measurements were performed in triplicate (n = 3), and data are reported as mean ± standard deviation unless otherwise stated.

## Data Availability

SAXS data have been deposited into SASBDB (Small Angle Scattering Biological DataBase) with accession codes: SASDZK8 (SmVAL13 alone), SASDZR8 (pAb anti SmVAL13 alone), SASDZL8 (SmVAL13 in complex with cognate pAb in ratio 2:1, 2^nd^ peak), SASDZM8 (SmVAL13 in complex with cognate pAb in ratio 2:1, 1^st^ peak), SASDZN8 (SmVAL13 in complex with its cognate pAb in ratio 1:1 (1^st^ peak), SASDZP8 (SmVAL13 in complex with its cognate pAb in ratio 1:1, 2^nd^ peak), SASDZQ8 (SmVAL13 in complex with cognate pAb in ratio 0.5:1, 2^nd^ peak); SASDZA8 (Polyclonal antibody anti-SmVAL11), SASDZH8 (SmVAL11 full length), SASDZJ8 (SmVAL11 auto-cleaved domains), SASDZB8 (SmVAL11 in complex with cognate pAb, ratio 2:1, 2^nd^ peak), SASDZC8 (SmVAL11 in complex with cognate pAb, ratio 2:1, 1^st^ peak), SASDZD8 (SmVAL11 in complex with cognate pAb, ratio 1:1, 2^nd^ peak), SASDZE8 (SmVAL11 in complex with cognate pAb, ratio 1:1, 1^st^ peak), SASDZF8 (SmVAL11 in complex with cognate pAb, ratio 0.5:1, 2^nd^ peak), SASDZG8 (SmVAL11 in complex with cognate pAb, ratio 0.5:1, 1^st^ peak).

All other data and materials are available upon request and following a material transfer agreement.

## Acknowledgments and Funding

We acknowledge SOLEIL for provision of synchrotron radiation facilities (project 20231630) and we would like to thank Dr Aurelien Thureau for assistance in using the beamline “SWING”. We acknowledge BAG beamtime MX2695 for accession to ESRF BM29. We would like to thank Dr Simon Megy for critical revision of the manuscript.

This project has received financial support from the CNRS through the MITI interdisciplinary programs “DetectSchisto” (AEM). FI doctoral project is funded by the CNRS 80|PRIME thesis “DetectSchisto”. SMR acknowledges the University of Lancashire for doctoral studentship support of ANS; and the Royal Society of Chemistry Research Enablement Grant (E22-5899202825). IE was supported by grants from MUR - “Progetto Eccellenza 2023 – 2027”.

## Author contributions

The study was conceived by AEM; FI, GB, AEM performed the biochemical and biophysical analyses; IE and FI performed the bioinformatics analysis; AG performed the MS analysis; ANS and SMR performed the bioaffinity reagents synthesis and analysis; AEM and SMR supervised the work; AEM, IE, SMR were responsible for funding acquisition; FI and AEM drafted the original manuscript with inputs from all the authors; all the authors edited and approved the final version.

