## Supplemental Tables 1-2; Suppl Figure S1 to S5 for "Biophysical characterization of novel biomarkers and bioaffinity reagents (NanoMIPs): on the road for a low-cost diagnostic test for intestinal schistosomiasis"

### Supplementary Material

**Supplementary Table 1. BLI kinetics and steady-state parameters for the three SmVAL recognition systems.** Kinetic $K_{D2}$ and $k_{off2}$ (mean ± SD per sensor) derive from the resolved second component of a 2:1 heterogeneous-ligand fit (retained cycles: $R^2 \geq 0.95$, $k_{off2} \geq 1 \times 10^{-5}$ $s^{-1}$). $K_{D1}$ was unresolved for the nanoMIP and fell below the instrument's resolution limit for both antibodies (< 1 pM). Steady-state $K_D$ (value ± fitting error) is from an independent Langmuir fit and is not directly comparable to the kinetic $K_{D2}$. t1B3 excluded from the steady-state group mean (poor isotherm fit, $R^2 = 0.87$).

| Group | Sensor | n | Kinetic $K_{D1}$ | Kinetic $K_{D2}$ (nM) | $k_{off2}$ ($s^{-1}$) | Kinetic $R^2$ | Steady-state $K_D$ (nM) | Steady-state $R^2$ |
|---|---|---|---|---|---|---|---|---|
| **Ab anti-SmVAL13** | t1E3 | 2 | < 1 pM | 91 ± 9 | $3.01\times10^{-4}$ | 0.99 | 500.0 ± 42.0 | 0.99 |
| | t1F3 | 4 | < 1 pM | 106 ± 62 | $2.79\times10^{-4}$ | 0.99 | 480.0 ± 37.0 | 0.99 |
| | t1G3 | 4 | < 1 pM | 106 ± 67 | $2.86\times10^{-4}$ | 0.99 | 390.0 ± 22.0 | 0.99 |
| | **Group mean** | | — | **103 ± 53** | **$2.9\times10^{-4}$ ± $0.5\times10^{-4}$** | **0.99** | **456.7 ± 58.6** | **0.99** |
| **Ab anti-SmVAL11** | t1A3 | 4 | < 1 pM | 986 ± 645 | $1.52\times10^{-3}$ | 0.98 | 85.0 ± 8.1 | 0.99 |
| | t1B3 | 2 | < 1 pM | 1825 ± 1411 | $2.33\times10^{-3}$ | 0.98 | 68.0 ± 14.0 | 0.87 |
| | t1C3 | 3 | < 1 pM | 359 ± 124 | $1.08\times10^{-3}$ | 0.98 | 95.0 ± 16.0 | 0.93 |
| | **Group mean** | | — | **963 ± 855** | **$1.6\times10^{-3}$ ± $0.6\times10^{-3}$** | **0.98** | **90.0 ± 7.1** | **0.95** |
| **MIP:SmVAL13** | t1A1 (20 µg) | 5 | Not resolved | 106 ± 18 | $8.96\times10^{-3}$ | 0.99 | 54.0 ± 1.8 | 0.99 |
| | t1B1 (40 µg) | 5 | Not resolved | 89 ± 50 | $8.86\times10^{-3}$ | 0.99 | 13.0 ± 1.0 | 0.97 |
| | **Group mean** | | — | **97 ± 35** | **$8.9\times10^{-3}$ ± $1.6\times10^{-3}$** | **0.99** | **33.5 ± 29** | **0.98** |

**Supplementary Table 2.** Comparative costs for biologicals in order to build a disposable portative diagnostic tests in large scale. Commercial point of care tests imply the immobilization of 50 ng of Ab or nanoMIP per test, therefore the cost of biologicals is 55-70 GBP per Ab-based sensor and 3 GBP /test based on nanoMIP.

| **Cost Category** | **NanoMIP Cost per mg (£)** (Reddy et al., 2024) | **Polyclonal Antibody Cost per mg (£)** | **References** |
|---|---|---|---|
| Materials | 0.29 | 2.5-10 | (Farid, 2007) |
| Labour / Specialists | 13.00 | 150-250 | (Klutz et al., 2016) |
| Energy & Facilities | 0.12 | 20-40 | (Farid, 2007) |
| Purification | 0 | 100-180 | (Xenopoulos, 2015) |
| Quality Control / Regulatory | 13.00 | 80-180 | |
| Target molecule (SmVAL13) | 20.9 (0.6 mg used to produce nanoMIP) | 750 (2.5 mg used to produce antibody) | |
| **Total Manufacturing Cost** | ~60.00 GBP | **1,100-1,410 GBP** | |

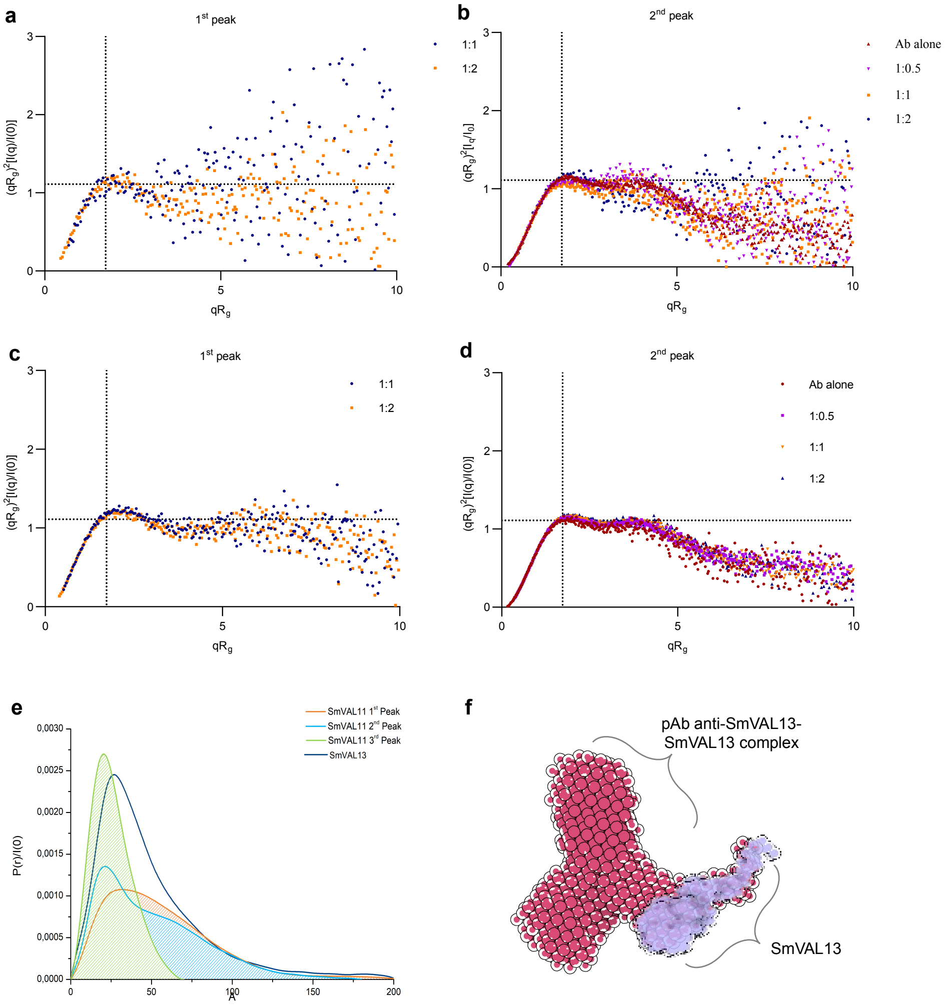


**Supplementary Figure S1|** Normalized Kratky plot of the 1st **(a)** and 2nd **(b)** eluted peak of pAb Anti-SmVAL13: SmVAL13 in different ratios; normalized Kratky plot of the 1st **(c)** and 2nd **(d)** eluted peak of pAb Anti-SmVAL11: SmVAL11 in different ratios. **(e)** Pr functions of the deconvoluted peaks. **(f)** *Ab initio* model of the complex Ab:SmVAL13, calculated with DAMMIF and averaged through DAMAVER, superposed with Coot [Casañal et al., 2020] to the *ab initio* model of SmVAL13 alone, calculated with DAMMIF and averaged through DAMAVER [Manalastas-Cantos et al., 2021].

```
CLUSTAL omega alignment

SmVAL11-N    -------------------------MSRLDDFREDCFRAHNEKRLLH------GVC------ALRHSLALDKTAQDWAEALLSEDGIKNSPLSSRG-E-VGESISV
SmVal11-C    -------------------------SKKLQESIHSVVQLHNQYRSQH------GSN------PLVLDQNLSNMAQQWADHLLQQSHLSNSGYVYRGMK-VGENLGS
SmVAL13      --------------------------MVDEQLNHDALNEHNRLRALH------GCP------PLKYDRRLAREAQAWAENLARLKIMKHSIC----DE-YGENLAS
SmVAL4       -MFKIVLVS---C--LLFLFTSLYVETKLSEGQRAIYNFHKKVRKDVKNCRIPGQPPAKNLTKLKWNKLLANKAKQQAKRCKYDSNDPNDFIIGDFES-IGQNLAD
4AIW.A       -----------------------MGKSASKQFHNEVLKAHNEYRQKH------G-VP-----PLKLXKNLNREAQQYSEALASTRILKHSPESSRGQC--GENLAW
5ETE.A       --------------------SASSSDSDLSDFASSVLAEHNKKRALHK------DTP-----ALSWSDTLASYAQDYADNYDCSGTLTHSGG------PYGENLAL
1CFE.A       -----------------------------QNSPQDYLAVHNDARAQV------G-VG-----PMSWDANLASRAQNYANSRAGDCNLIHSG--------AGENLAK
5V50.A       -----------------PAAELEARQFDPDSFKNKWLELHNNERTTRQ-------LD-----SLEWDGDLAWKAQQVATQCNVDNPQL-----------WGDNGAS
4TPV.A       -----------------DYSLCQQREKLDDDMREMFTELHNGYRAAFARNYKTSKMR-----TMVYDCTLEEKAYKSAEKCS----EEPSSE--------EENVDV
                                                    **  *                     ..  *    .              :.         .::.

SmVAL11-N    RTSTGTHVDMQGHEVVNTWHSDAENYNYENG----KGPAGNFTQLVWSSTREVGFGKACG----------PGKCVVVAHYRPPGNVLGRYLENVFRPKKSVKDVKQ
SmVal11-C    RWSNG-PMELNCKDLIEHWYQESGKYKFNSEPDS-IQGIGNFTQIVWSSSERIGVGIAIQSYKSGEDLHKDSKMILVCLYHPPGNVISQFQNNVKKAIK-------
SmVAL13      AQSTG-KAEMTGARATRNWYDEIHYHNFNKQF---QSQSGHFTQLIWKNTSKAGFG--IQHSV------DGHHVFIVGRYEPPGNVNGQFLENVPPPIHGQSTPKS
SmVAL4       YPTIE--------GAMKDWLEEYKNYNFEKNQCNG--DCKNYKQMVWNTTEEIGCGYEKC----GKNY------LIVCNYAP-GDSE-DRPYEAKPESKCNKSET-
4AIW.A       ASYDQ-----TGKEVADRWYSEIKNYNFQQPGFTS--GTGHFTAMVWKNTKKMGVGKASA-----SD----GSSFVVARYFPAGNVVNEGFFEENVLPPKK-----
5ETE.A       G---Y-----DGPAAVDAWYNEISNYDFSNPGFSS--NTGHFTQVVWKSTTQVGCGIKTC-----GG---AWGDYVICSYDPAGNY--EGEYADNVEPLA------
1CFE.A       GGGDF-----TGRAAVQLWVSERPSYNYATNQCVGGKKCRHYTQVVWRNSVRLGCGRARC-----NN----GWWFISCNYDPVGNWIGQRPY--------------
5V50.A       -FNIG-R--YTKEQAFAEWTATSGSFP------DD--RSIPWQRIVANSAQKVGCGEATC-----VLEGDMAYTVNVCYYDPPLSDYYTNAGDNVRVPSLALLNLD
4TPV.A       -FSAAT--LNIPLEAGNSWWSEIFELRGKVYNKNG--KTSNIANMVWDSHDKLGCAVVDC-----S-----GKTHVVCQYGPEAKGDGKTIYEEGAPCSRCSDYGA
                           :   *:.:                   :...::.   . *::                       * *  .

SmVAL11-N    PIRNTF-ALNNDTPKTVITETLT---ESDGK-----------------------QYSVRREISDLTDDKGKTRRCINEVYTDACKEQKKAKSSISPDGHLVDN
SmVal11-C    -------------------------------------------------------------------------------------------------------
SmVAL13      KVPSYK-HNEQNGPRRTYQDELVIVRETDRKDHNGSN-HIT-LIDSSKRSRSDETIPNKNEVRIIRAEKKRQRRCAKRCSIM---------------------
SmVAL4       -------------------------------------------------------------------------------------------------------
4AIW.A       -------------------------------------------------------------------------------------------------------
5ETE.A       -------------------------------------------------------------------------------------------------------
1CFE.A       -------------------------------------------------------------------------------------------------------
5V50.A       LTTPFCNLPVNGSNDLDIRE-----------------------------------------------------------------------------------
4TPV.A       GVTCDDDWQNLLCIGHHHH------------------------------------------------------------------------------------
```

**Supplementary Figure S2|** Multiple alignment with Clustal Omega [Madeira et al., 2024] of SmVALs and templates used for homology modeling. PDB IDs: 4AIW.A – human GAPR1, 5ETE.A, 1CFE.A, 5V50.A – *Moniliophthora perniciosa* MpPR-1i, 4TPV.A - *Ancylostoma caninum* Hookworm Platelet Inhibitor. The sequences of SmVAL11-N, SmVAL11-C and SmVAL13 have been colored following the secondary structure and epitope mapping predicted by BepiPred [Clifford et al., 2022], (grey: disordered; orange: coil; red: alpha-helix; blue: beta-strand).

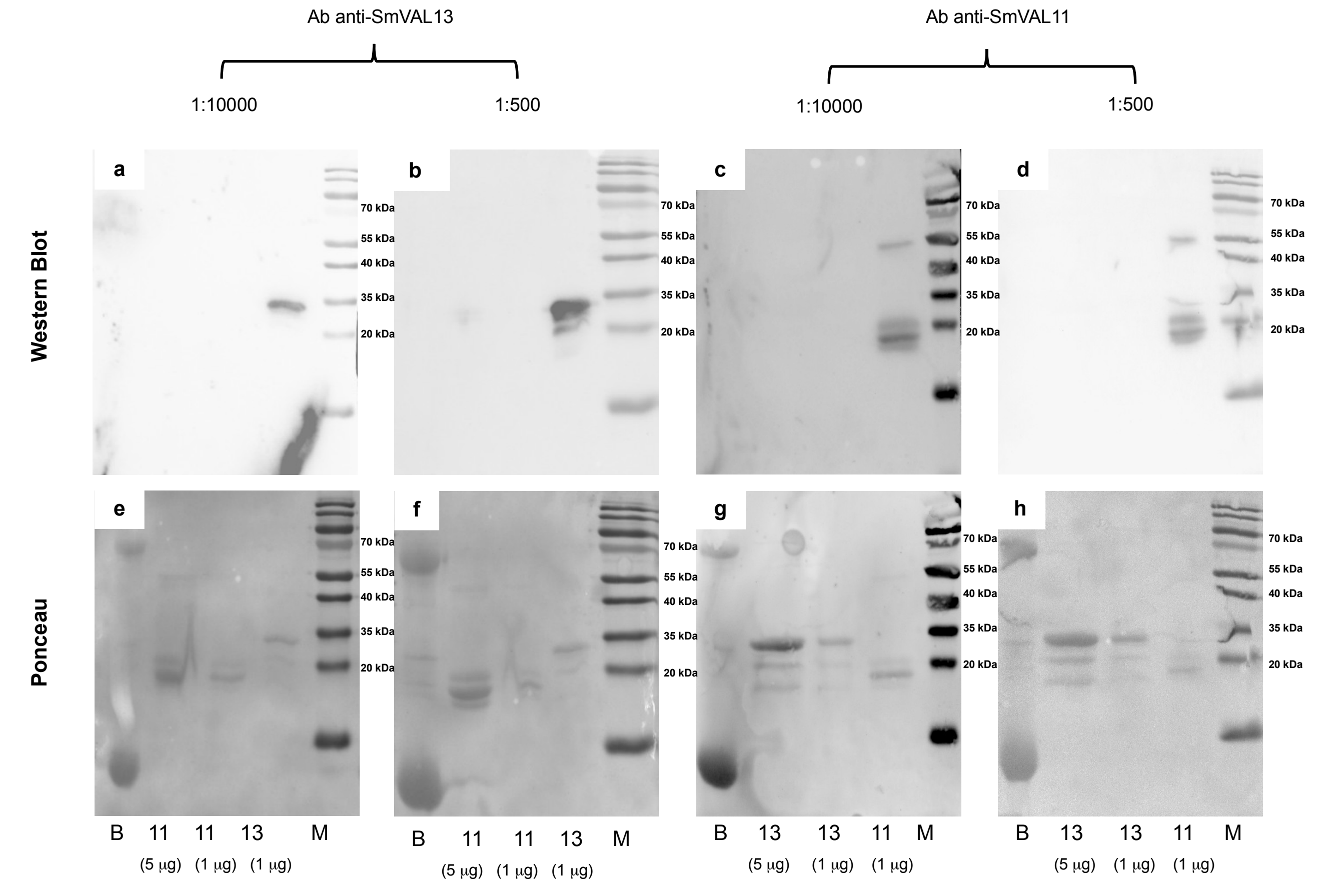


**Supplementary Figure S3| Specificity assessment of anti-SmVAL13 and anti-SmVAL11 polyclonal antibodies by Western blot.** Western blot analysis of recombinant SmVAL11 and SmVAL13 proteins using pAb anti-SmVAL13 (a,b) and pAb anti-SmVAL11 (c,d) at two different dilutions: 1:10,000 (a,c) and 1:500 (b,d). Corresponding Ponceau S staining of each membrane is shown below as a loading and transfer control (e–h).

Lane **B**: whole blood (about 20 µg) from non infected human; lanes **11**: recombinant SmVAL11 (5 µg or 1 µg, as indicated); lanes **13**: recombinant SmVAL13 (5 µg or 1 µg, as indicated). **M**: Molecular weight markers (kDa). The anti-SmVAL13 antibody specifically recognizes SmVAL13 with no detectable cross-reactivity toward SmVAL11 or blood proteins, whereas the anti-SmVAL11 antibody specifically detects SmVAL11 without cross-reactivity toward SmVAL13.

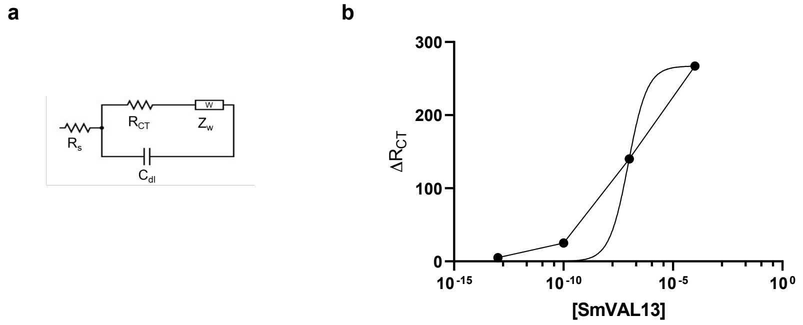


**Supplementary Figure S4| (a)** Randles–Warburg equivalent circuit used to fit the EIS data, where $R_S$ represents the solution resistance, $R_{CT}$ the charge-transfer resistance, $C_{dl}$ the double-layer capacitance, and $Z_W$ the Warburg diffusion impedance. **(b)** Change in charge-transfer resistance (ΔRct) measured by EIS as a function of SmVAL13 concentration, fitted with a Langmuir Isotherm.

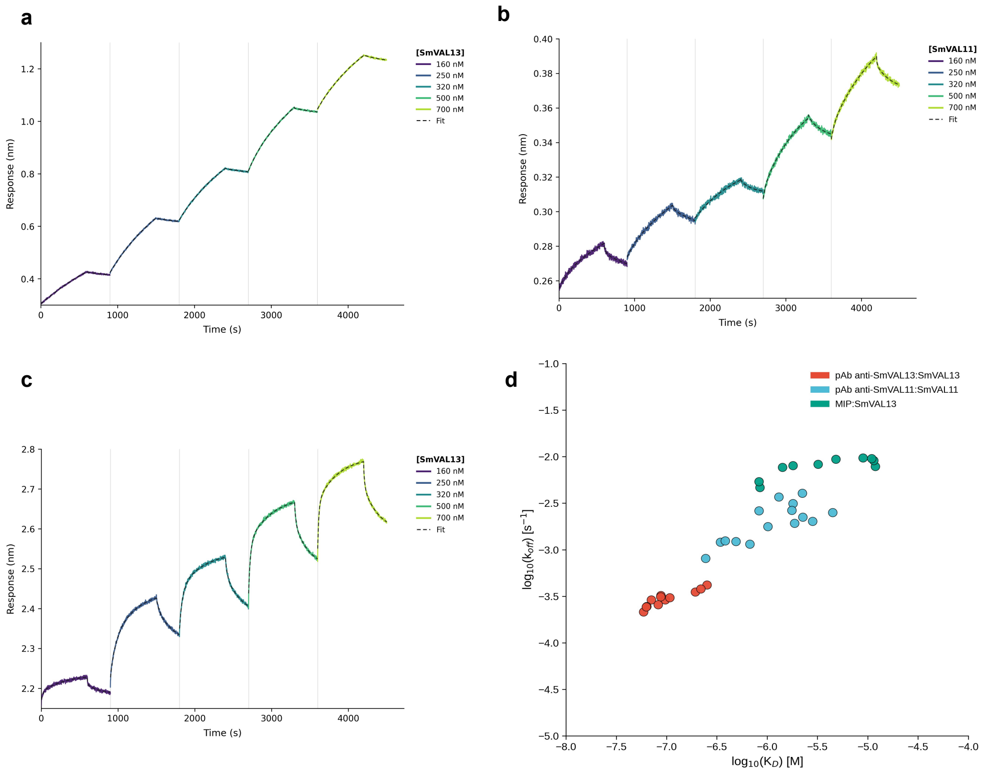


**Supplementary Figure S5|** Single-cycle BLI sensorgrams of **(a)** immobilised pAb-antiVAL13 vs SmVAL13, **(b)** immobilized pAb-antiVAL11 vs SmVAL11 and **(c)** immobilized nanoMIP anti-VAL13 vs SmVAL13. Increasing analyte concentrations (160–700 nM) were tested sequentially without regeneration; vertical lines mark each association-dissociation step. (**d**) Kinetic landscape of SmVAL11/13 recognition by cognate Ab and MIP sensors. Scatter plot of $\log_{10}(k_{off})$ as a function of $\log_{10}(K_D)$, derived from the second binding population of a 2:1 heterogeneous ligand model fitted to BLI single-cycle kinetics data. Each point represents an individual analyte concentration tested on a given sensor. Colors indicate the antibody/sensor–antigen pair: pAb anti-SmVAL13:SmVAL13 (red, n = 12), pAb anti-SmVAL11:SmVAL11 (cyan, n = 15), and nanoMIP:SmVAL13 (green, n = 10).
